# Early Prediction of Antigenic Transitions for Influenza A H3N2

**DOI:** 10.1101/558577

**Authors:** Lauren A Castro, Trevor Bedford, Lauren Ancel Meyers

## Abstract

Influenza A/H3N2 is a rapidly evolving virus which experiences major antigenic transitions every two to eight years. Anticipating the timing and outcome of transitions is critical to developing effective seasonal influenza vaccines. Using simulations from a published phylodynamic model of influenza transmission, we identified indicators of future evolutionary success for an emerging antigenic cluster. The eventual fate of a new cluster depends on its initial epidemiological growth rate––which is a function of mutational load and population susceptibility to the cluster––along with the variance in growth rate across co-circulating viruses. Logistic regression can predict whether a cluster at 5% relative frequency will eventually succeed with ∼80% sensitivity, providing up to eight months advance warning. As a cluster expands, the predictions improve while the lead-time for vaccine development and other interventions decreases. By focusing surveillance efforts on estimating population-wide susceptibility to emerging viruses, we can better anticipate major antigenic transitions.

## INTRODUCTION

Seasonal influenza A/H3N2 causes significant annual morbidity and mortality worldwide, as well as severe economic losses [1]. In the United States, the 2017-2018 season was unusually long and severe, lasting over 16 weeks and causing over 900,000 hospitalizations and 80,000 fatalities, including 183 pediatric deaths [2–4]. The global health community continually tracks H3N2 and annually updates the H3N2 component of the seasonal influenza vaccine. However, annual influenza epidemics continue to impart a significant public health burden. The rapid antigenic evolution of the influenza virus via mutations in hemagglutinin (HA) glycoproteins and neuraminidase (NA) enzymes [5,6], and logistical requirement of selecting vaccine strains almost a year prior to the flu season pose a significant challenge. Vaccines target the antigen-binding regions of dominant influenza subtypes. While a particular subtype may circulate for a few years, strong positive selection for new antigenic variants will eventually produce antigenic drift [7–9], rendering a vaccine less effective if new mutations in the antigen-binding regions are not included in vaccine chosen strains [10,11]. The typical reign of a dominant subtype ranges from two to eight years [12,13]. From 2004 to 2018, seasonal influenza vaccines have had an estimated average efficacy of 40.56% against all influenza strains included in the vaccine [14].

The World Health Organization’s Influenza Surveillance and Response System (GISRS) coordinates influenza surveillance efforts to survey and characterize the diversity of influenza viruses circulating in humans. Viral samples are rapidly analyzed via sequencing of HA and NA genes, serologic assays, and other laboratory tests to identify newly emerging antigenic clusters. Within the past decade, the number of complete HA gene sequences in the GISAID EpiFlu [15,16] database has increased tenfold, from fewer than 1,000 in 2010 to over 10,000 in 2017 [17]. Molecular data at high spatiotemporal resolution could potentially revolutionize influenza prediction. However, the research and public health communities have just begun to determine effective strategies for extracting and integrating useful information into the vaccine selection process.

Phylodynamic models describe the interaction between the epidemiological and evolutionary processes of a pathogen [18]. The availability of molecular data coupled with the recent development of detailed, data-driven phylodynamic models has galvanized the new field of viral predictive modeling [19–22]. These models aim to predict the future prevalence of specific viral subtypes based on past and present molecular data. For example, one approach generates one-year ahead forecasts of clade frequency using a fitness model parameterized by the number of antigenic and genetic mutations that dictate the virus’ antigenicity and stability respectively [23]. Another method maps antigenic distance from hemagglutination inhibition (HI) assay data onto an HA genealogy to determine whether the changes in antigenicity among high-growth clades necessitate a vaccine composition update [24]. A third model predicts which clade will be the progenitor lineage of the subsequent influenza season by estimating fitness using a growth rate measure derived from topological features of the HA genealogy [25]. All three approaches have been tested on historical predictions. Łuksza’s & Lässig’s model [23] predicted positive growth for 93% of clades that increased in frequency over one year. Steinbruk *et al.* [24] predicted the predominant HA allele over nine influenza seasons with an accuracy of 78%. Both Łuksza’s & Lässig’s [23] and Neher *et al.* [25] model predictions of progenitor strains to the next season’s performed similarly. Since 2015, both these models have been used to provide recommendations on vaccine composition for the upcoming influenza seasons [26–28].

Taken together, this body of work points to the promise of predictive evolutionary models. Phylodynamic simulation models provide a complementary window into the molecular evolution of emerging viruses. By observing influenza evolution *in silico*, we can take rigorous experimental approach to test hypotheses about early indicators of cluster [29,30] success and design surveillance strategies to inform vaccine strain selection. Here, we simulate decades of H3N2 phylodynamics using a published model [31,32] and analyze the simulated data to identify early predictors of a cluster’s evolutionary fate. Viral growth rates––both for an emerging cluster and its competitors––are the most robust predictors of future ascents. When a new antigenic cluster first appears at low frequency (e.g., 1% of sampled viruses), our models can predict whether it will eventually rise to dominance (e.g., maintain a relative frequency greater than 20% of sampled viruses for at least 45 days) with reasonable confidence and advanced warning. To translate these findings into actionable guidance for global influenza surveillance, we also evaluate proxy indicators that can be readily estimated from current data, quantify limits in the accuracy, precision and timeliness of predictions, and construct models to predict future frequencies of emerging clusters.

## METHODS

### Simulation Model

We implemented a published stochastic individual-based susceptible-infected (SI) phylodynamic model of influenza A/H3N2 [31,32] to repeatedly simulate 30 years of transmission in a constant population of 40 million hosts with birth and death dynamics (Fig. 1A). In brief, each individual host is characterized by its infection status––susceptible or infected––and a history of prior viral infections. Viruses are defined by a discrete antigenic phenotype, which determines the degree of immune escape from other phenotypes, and a deleterious genetic mutation load (*k*) which affects the virus’ transmissibility. Antigenic mutations occur stochastically and confer advanced antigenicity to the virus. The probability that a given virus will infect a given host is determined by how similar the antigenic phenotype of the challenging virus is to the antigenic phenotype of the host’s most related previous infection. This probability, or degree of immune escape, is tracked through the simulation by the evolutionary history of clusters (parent-child relationships). Antigenic and deleterious non-antigenic mutations occur only during transmission events; the model assumes that viruses within a single individual host are genotypically homogeneous. The model also assumes no co-infection, no seasonal forcing [33,34], and no short-term immunity that would broadly prevent reinfection after recovering from infection. We used the baseline parameters chosen in Koelle & Rasmussen [31] based on empirical epidemiological and virological estimates [35–38].

**Figure 1:**
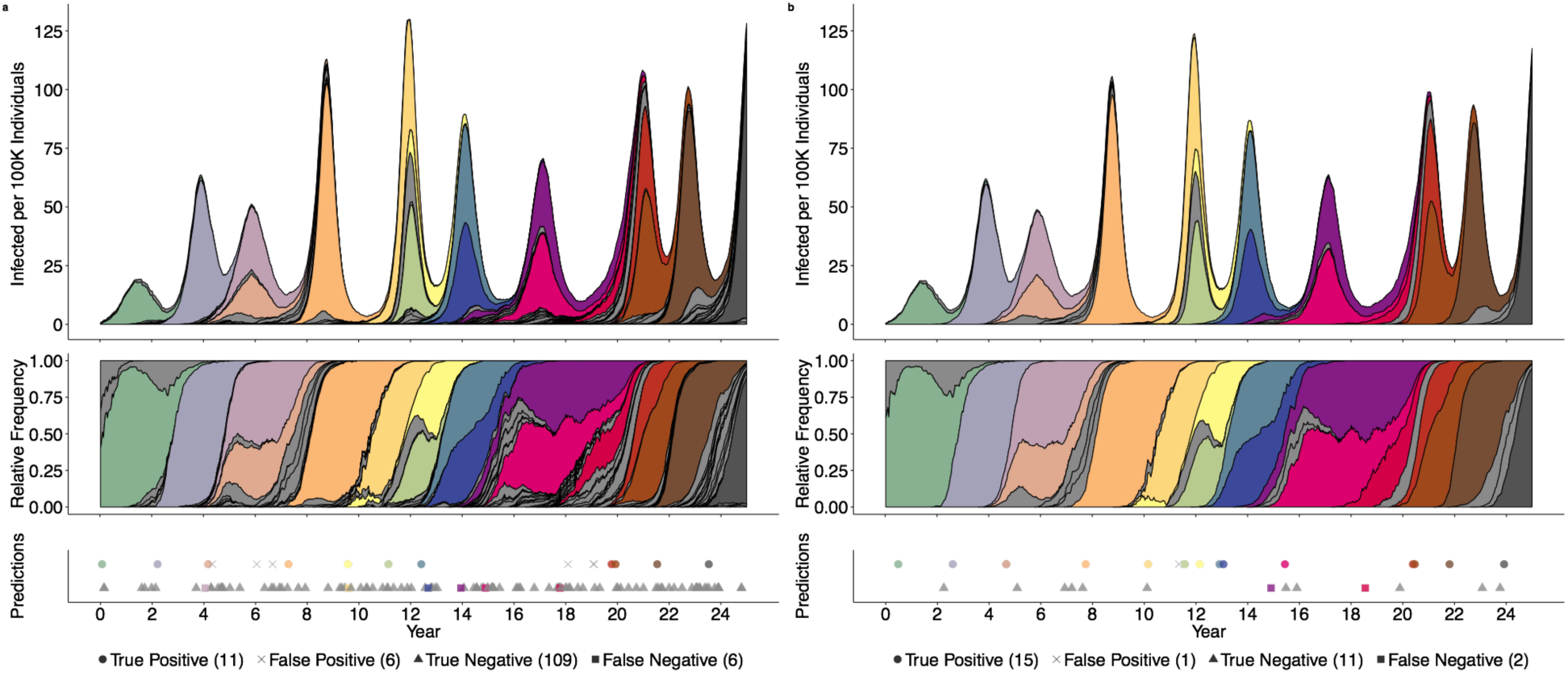
Out-of-sample predictions of antigenic cluster evolutionary success at relative frequency thresholds of 1% (a) and 10% (b). Grey shading indicates clusters that surpass the surveillance threshold, but do not establish. Other colors correspond to distinct antigenic clusters that eventually establish. The top time series graphs depict the absolute prevalence of antigenic clusters; the middle graphs give their relative frequencies. The bottom panels indicate the timing and accuracy of out-of-sample predictions based on the optimized model for each surveillance threshold. The top row of symbols indicate clusters predicted to succeed, with true positives indicated by circles and false positives indicated by crosses; the bottom row indicates clusters predicted to circulate only transiently, true negatives indicated by triangles and false negatives indicated by squares. The number of predictions in each category is provided in the legend.

### Simulated Data

We ran 100 replicate simulations and selected a subset that produced realistic global influenza dynamics. Specifically, we excluded 38 simulations in which endemic transmission died out prior to the 30 years. We treated the first five years of each simulation as burn-in periods. In total, we analyzed 1550 years of simulated influenza transmission and evolutionary dynamics.

Throughout each simulation, we tracked 23 metrics reflecting the epidemiological state of the host population (i.e., number of susceptible and infected individuals) and evolutionary state of the viral population (Table S1) at 14 day intervals. When possible, we monitored these quantities for both individual antigenic clusters and the entire viral population, and then calculated their ratio. For example, we monitored the average number of deleterious mutations within each antigenic cluster and across all viruses, as well as the *relative* mutational load of each cluster with respect to the entire viral population. Henceforth, we refer to the metrics as *candidate predictors*.

### Classifying Evolutionary Outcomes

We classified each novel antigenic cluster in each simulation into one of three categories: (1) rapidly eliminated clusters that never reach 1% relative frequency in the population, (2) transient clusters that surpass 1% relative frequency but do not qualify as established clusters, and (3) established clusters that circulate above 20% relative frequency for at least 45 days. With this criteria, transient and established clusters constituted on average 81% of the infections at any point in time (Figs S1-2).

### Predictive Models

Restricting our analysis to transient and established clusters, we used generalized linear modeling to identify important early predictors of evolutionary fate. For each antigenic cluster, we predicted its evolutionary future (i.e., whether it ultimately becomes established) at specified surveillance thresholds, such as 5% relative frequency. Specifically, we recorded all candidate epidemiological and evolutionary predictors at the moment each cluster crossed the threshold. We analyzed all ten surveillance thresholds ranging from 1% to 10% at 1% increments.

For each surveillance threshold, we centered and scaled candidate predictors and removed collinear factors. Using five-fold cross validation, we partitioned the data into five subsets, keeping data from individual simulations in the same subsets. We fit mixed-effects logistic regression models using four subsets for training and controlling for differences between independent simulations. Predictors were added sequentially based on which term most significantly lowered the average Akaike Information Criterion of the five training folds.

We evaluated model performance by predicting the evolutionary outcomes of clusters in the held-out test subset. We calculated three metrics: the area under the receiver operating curve (AUC), the sensitivity (the proportion of all positives predicted as positive), and the positive predictive value (the proportion of true positives of all predicted positives). The model predicts the probability that a cluster will establish. To translate these outputs into discrete binary predictions of future success, we applied a probability threshold which maximized the F1 score [39], which is the harmonic average of a model’s positive predictive value and sensitivity (Table S2). When we included historical data of candidate predictors, i.e the value of a candidate predictor at an earlier surveillance threshold, model performance did not have a significant difference (Fig. S3).

We also considered an opportunistic sampling regime, where samples are tested as they arise regardless of their relative frequency. We fit models aimed at two prediction targets: (1) the evolutionary success of a cluster sampled at an arbitrary relative frequency and (2) the frequency of a cluster up to twelve months into the future. We built models based on data sampled from ten random time points in each of the 62 25-year simulations. We considered all clusters present above 1% relative frequency but not yet established as a dominant cluster. The frequency of a cluster at the time of sampling was included as an additional predictor. To predict the frequency of an antigenic cluster *X* months into the future, we fit a two-part model that first predicted whether the cluster would be present at the specified date, and, if so, then estimated the frequency of the cluster at that date. We used forward variable selection and cross validation model, as described above. We used the R statistical language version 3.3.2 [40] for all analyses, and the *afex* package for generalized linear models [41].

### Candidate predictors

#### Reproductive Rates

In our simulated data, we can calculate the instantaneous reproductive rate for particular clusters and the entire viral population. As described in Koelle & Rasmussen (2015), the reproductive rate of a virus *v* is given by:

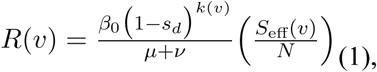

where *β*_0_ is the inherent transmissibility, *s*_*d*_ is the fitness effect for each of the virus’ *k* (*v*) deleterious mutations, *μ* and *ν* are the per capita daily death and recovery rates, respectively, and *N* is the host population size. We assume that *β*_0_, *s*_*d*_, *μ* and *ν* are constant across all viruses. *S*_eff_ (*v*) denotes the population-wide susceptibility to the virus accounting for cross-immunity from prior infections, herein referred to as the effective susceptibility, and the population level effective susceptibility is estimated for a virus as:

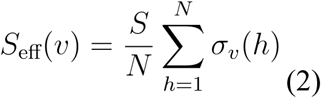

where *σ*_*v*_ (*h*) is the immunity of host *h* towards virus *v* based on the antigenic similarity between *v* and the virus in host *h*’s infection history most antigenically similar to virus *v*. A *σ*_*v*_ (*h*)=1 indicates full susceptibility, while *σ*_*v*_ (*h*) =0 indicates complete immunity.

The growth rate of an antigenic cluster is then the average over all viruses in that cluster, given by:

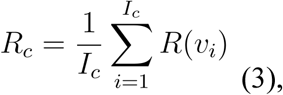

where *I*_*c*_ is the number of hosts infected by a virus from cluster *c* and *v*_*i*_ is the virus infecting host *i*. Likewise the population-wide average (⟨*R* ⟩) and variance (var (*R*)) in *R* are computed across all current infections, and the relative reproductive rate of a cluster is given by *R*_*c*_/⟨*R*⟩

#### Practical approximations

Eqs. (1-3) are not easily calculated from current surveillance data. Therefore, we considered two proxy measures of viral growth rates and two proxy measures of viral competition. We first choose two surveillance thresholds, for example, 6% and 10%. When the relatively frequency of a cluster crosses the second threshold, we calculate both the *time elapsed* since it crossed the first threshold and the *relative fold change*, as given by:

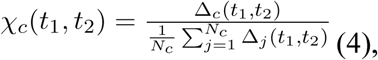

where *t* (1) and *t*(2) are the times at which cluster crossed the first and second threshold, respectively, Δ_*c*_ (*s,t*) is its relative frequency at time *t* divided by its relative frequency at time *s* and *N*_*c*_ is the number of distinct clusters present at both time *t*(1) and *t*(2). For the competition proxy measures, we calculate the the variance in *𝒳*_*c*_(*t*_1_,*t*_2_) and the *N*_*c*_ where Δ_*c*_ (*s,t*) > 1.

We evaluate the performance of these approximations by comparing logistic regression models that predict whether a cluster will establish from either the true *R*_*c*_/⟨*R*⟩ at the 10% surveillance threshold, the relative fold change between the 6% and 10%, or time elapsed between reaching the 6% and 10% thresholds. As before, we evaluated model performance based on AUC, positive predictive value, and sensitivity.

## RESULTS

Our simulations roughly reproduce the global epidemiological and evolutionary dynamics of H3N2 influenza over a 25 year period. Without seasonal forcing, prevalence rises and falls, peaking every 3.2 years on average (s.d. = 1.6). These dynamics reflect the turnover and competition of antigenic clusters. The median of the most recent common ancestor (TMRCA) in our simulations was 5.9 years (IQR 4.62 - 7.9), which is higher than empirical estimates of 3.89 years [42]. The median life span of established clusters was 1128 days (s.d. = 480), corresponding to roughly 3.5 years. However, the annual incidence of influenza in our model (4.0%, 95%CI 0.37-9.7%) was lower than empirical annual incidence estimates of 9–15% [42]. Given the model only simulates the transmission of H3N2 and not all circulating influenza types, our annual incidence is comparable to empirical estimates [43].

We assume that clusters become detectable once they cross a relative frequency threshold of 1% and are fully established if they maintain a relative frequency above 20% for at least 45 weeks. In our simulations, 2% of the approximately 200 novel antigenic clusters per year overcome early stochastic loss to reach detectable levels. As the relative frequency of a newly emerging cluster increases, the probability that the cluster will ultimately establish also increases. There is an inverse relationship between the number of clusters that reach a threshold and the probability of future success. For example, far fewer clusters reach a relative frequency of 10% than 1%. If a cluster succeeds in reaching relative frequency thresholds of 1%, 6%, and 10%, its probability of establishing increases from 13% to 50% to 67% (Fig. S2).

Our model classifies clusters as either *positives* that are likely to establish or *negatives* that are expected to circulate only transiently. As we increase the surveillance threshold, the fraction of successful clusters that are misclassified as negatives decreases. In a representative out-of-sample 25-year simulation, 17 of 132 detectable clusters eventually rose to dominance (Fig. 1). Of these, 65% and 88% were correctly predicted when they reached the 1% and 10% surveillance threshold, respectively. The number of true negative events decreased considerably, from 109 at the 1% surveillance threshold to only 11 at the 10% surveillance threshold, while the other types of events held relatively constant.

Across all surveillance thresholds, the first four predictors chosen through forward model selection are the relative growth rate of the focal cluster (*R*_*c*_/⟨*R*⟩), the background variance (var (*R*)) and mean (⟨*R*⟩) of viral growth rates, and the relative deleterious mutational load of the focal cluster (*k*_*c*_/⟨*k*⟩). Population-level epidemiological quantities were only selected for models at low surveillance thresholds (2-4%); in these models, overall prevalence had a slightly negative correlation with future viral success (Table 1). The median number of predictors chosen was 6.5, with a range of 5 to 7. The best fit models are described in Tables S2.

**Table 1:**
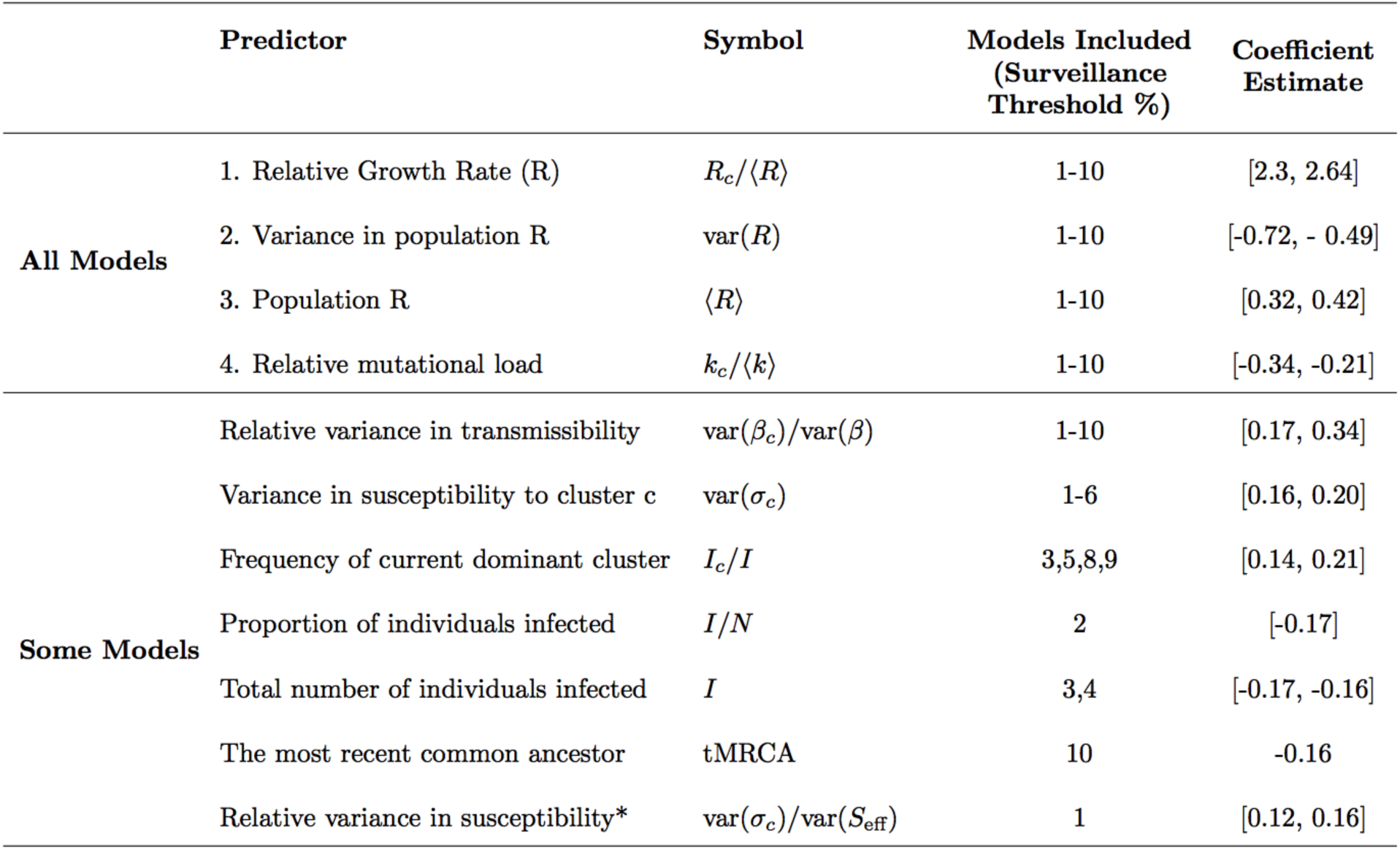
Predictors selected by five-fold cross validation and forward selection. The top four variables were selected in the identical order (as listed) across all surveillance threshold models. The fifth predictor, relative variance in transmissibility, was included in all models, but not always as the fifth chosen. In the formulas, refers to cluster-level quantities. The rightmost column gives the full range of fitted coefficients (log-odds) across all models based on across the five-fold cross validation for each surveillance thresholds’ final model. ^*^ var (*σ*_*c*_) was calculated across all hosts; var (*S*_eff_) was calculated across only infected hosts. *I*= number of infected hosts, *N*= total number of hosts, *σ*_*c*_ = effective susceptibility to infection by cluster c, *β*^*k*^ = the transmission rate of the virus carrying *k* deleterious mutations. Formulas to calculate each quantity are in Table S1.

We examine the dynamics of the top two predictors. As newly emerging clusters rise in relative frequency from 1% to 10%, their relative growth rate declines towards one. That is, they approach the population average fitness (Fig. 2). The relative growth rate is significantly higher for clusters that will eventually establish than those will burn out, with the separation between the two groups increasing as the clusters ascend in frequency (Fig. 2a). This predictor is a composite quantity, estimated based on both mutational load and effective susceptibility. We compare these two quantities at two time points, when the clusters reach 1% and 10% frequencies. Mutational load increases and effective susceptibility decreases in ascending clusters, with more extreme changes occurring in clusters that ultimately fail to establish. We also measure the changes in these two quantities for the entire population, and find that the background mutational load remains relatively constant and background effective susceptibility increases slightly. The background effective susceptibility peaks when a new cluster begins to constitute a major proportion of the circulating types––at this point the immunity from previous infections is not strongly protective against the newly dominant cluster. The decline in cluster fitness likely stems from the accumulation of deleterious mutations and exhaustion of the susceptible population (Fig. 2B). While this occurs within both established and transient clusters, the mutational loads in established and transients increase by averages of 1.4 and 2.04 mutations, respectively (Wilcox, p < 2.2e-16).

**Figure 2:**
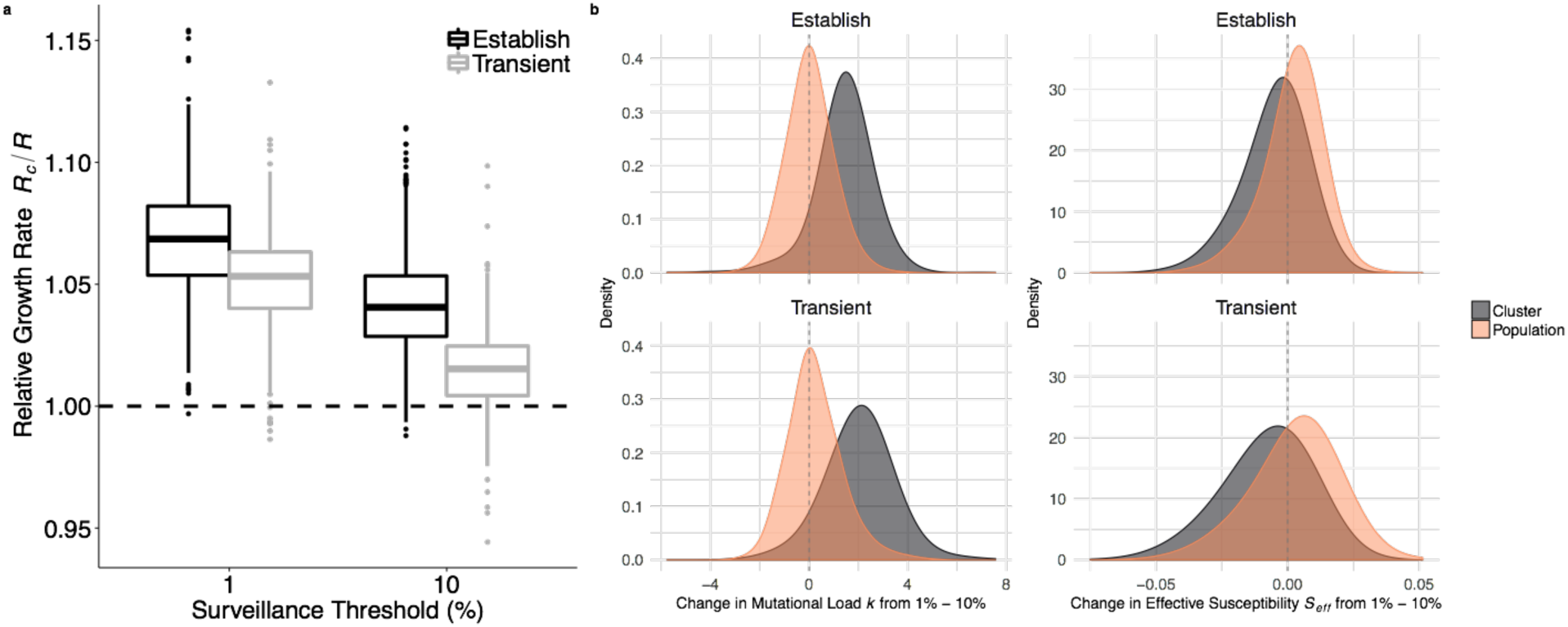
Relative growth rates predict future success. (a) Clusters that eventually establish have significantly higher *R*_c_/ ⟨*R* ⟩than those that fail to establish. As clusters increase in relative frequency from 1% to 10%, their generally declines but the distinction between future successes and future failures becomes more pronounced. (b) *R* is a composite value based on the mutational load and *S*_eff_. We compared the mutational load (left) and *S*_eff_ (right) of a cluster when it crossed the 1% and 10% thresholds by subtracting the former from the latter (orange distributions); we simultaneously calculated the difference in average mutational load and *S*_eff_ across the entire viral population (grey distributions). The top and bottom rows shows the distributions of change for clusters that establish and transiently circulate, respectively. The decrease in a cluster’s fitness advantage is driven by both increasing mutational load and a decreasing *S*_eff_. The background mutational load does not change noticeably, while the background *S*_eff_ increases slightly.

The background variance in viral growth rates, var(*R*), is the second most informative predictor. The lower the variance, the more likely a cluster is to establish. However, it is a weaker predictor than *R*_*c*_/ ⟨*R* ⟩ ’s; the estimated logit coefficient of the *R*_*c*_/ ⟨*R* ⟩ is approximately four times that of var(*R*) (Table 1). The var(*R*) tends to increase as a cluster expands from 1% to 10% relative frequency (Wilcox, p < 2.2e-16). This may stem from diverging fitnesses of the newly expanding cluster and the receding dominant cluster, which has likely accumulated a considerable deleterious load and burned through much of its susceptible host population. A higher var(*R*) decreases the probability of a cluster being successful, particularly when a cluster has only a modest growth rate. Clusters with high *R*_*c*_/ ⟨*R* ⟩ ’s are successful even when emerging in highly variant environments (Fig. 3A). High variance may reflect high levels of inter-viral competition. If we consider both transient and established clusters with similar *R*_*c*_/ ⟨*R* ⟩ (ranging from 1.025 to 1.03), successful clusters encounter significantly fewer co-circulating clusters, and the frequency of the resident dominant cluster is significantly higher (Fig. 3C). This may reflect suppression of competition by the dominant cluster, creating a vacuum for a moderately fit cluster to fill.

**Figure 3:**
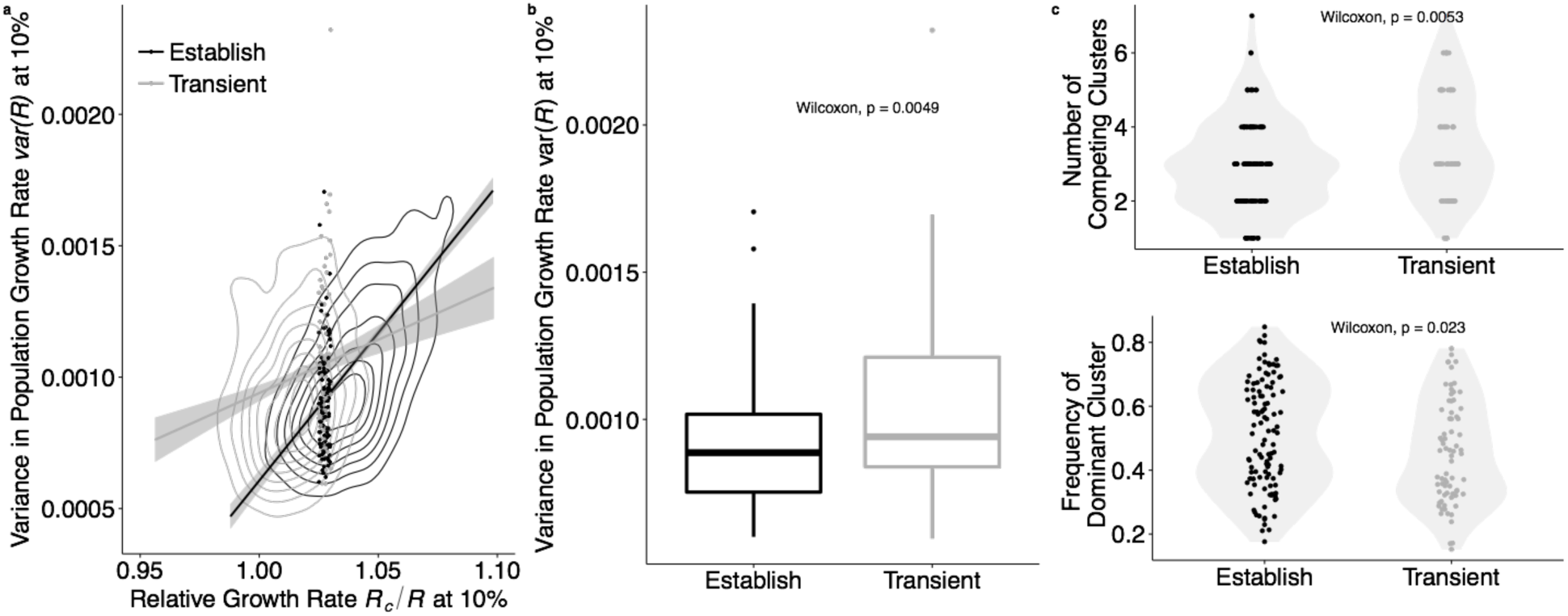
Viral competition predicts future success for clusters with borderline growth rates. (a) Clusters with only a slight *R*_*c*_/ ⟨*R* ⟩ advantage are more likely to establish if the background var(*R*) is low. Clusters with higher *R*_*c*_/ ⟨*R* ⟩ successfully regardless of .var (*R*) Contour lines indicate the density of values of and var(*R*). The lines represent the correlation between the variables for successful and transient clusters. (Success-black: r = 0.63, p < 2.2e-16; Transient-grey: r = 0.18, p <1.2e-06). The dots represent clusters with *R*_*c*_/ ⟨*R* ⟩ ’s between 1.025-1.030, a range within the individual distributions of *R*_*c*_/ ⟨*R* ⟩ for success and transient clusters do not statistically differ (Wilcox, p = 0.4551). For clusters falling within this ambiguous range of *R*_*c*_/ ⟨*R* ⟩, (b) var(*R*) is significantly higher in transient clusters than in established clusters, and (c) in comparison to transient clusters, successful clusters tend to face fewer co-circulating clusters (Wilcox, p = 0.0053), with the current dominant cluster at higher frequency (Wilcox, p = 0.023). Points represent the number of circulating clusters and the frequency of the dominant cluster; shading represents the kernel density estimation of the distribution of points. Across all graphs, values are calculated when the focal clusters reach a 10% surveillance threshold.

Using 6271 geographically diverse influenza A/H3N2 sequences sampled from 2006 and 2018, we assessed whether our predictive models can be directly applied to influenza surveillance efforts. Clusters were distinguished by single mutations to epitope sites on the HA1 sequence and successful clusters were those that reached a relative frequency of at least 20% for at least 45 days. Despite sparse sampling, the dynamics of antigenic transitions resemble those produced by our simulations (Fig. 4a). Over the 12-year period, dominant clusters circulated for an average of 2.25 years (sd. 1.17); 44 clusters reached a relative frequency of 10%; 18 of the 44 were eventually successful. For each emerging cluster, we calculated the relative number of epitope mutations by dividing the average number of epitope mutations in viruses within the cluster by the average number found in other co-circulating viruses. For clusters that reached 1% relative frequency, this quantity was less than one for clusters that eventually established (N=18) and greater than one for transient clusters that did not establish (N=1516); this difference is statistically significant (Wilcox, p = 0.0003) (Fig. 4b). This difference was not significantly different when measured when clusters reached the 5% relative frequency surveillance threshold. We also fit classifier models to the empirical data using proxies for fitness (e.g., fold change and growth rate between sequential sampling of a cluster) and competition, (e.g., the number of co-circulating clusters). While some of these factors are significant predictors of future evolutionary success, our best models had sensitivity and positive predictive values below 50%.

**Figure 4:**
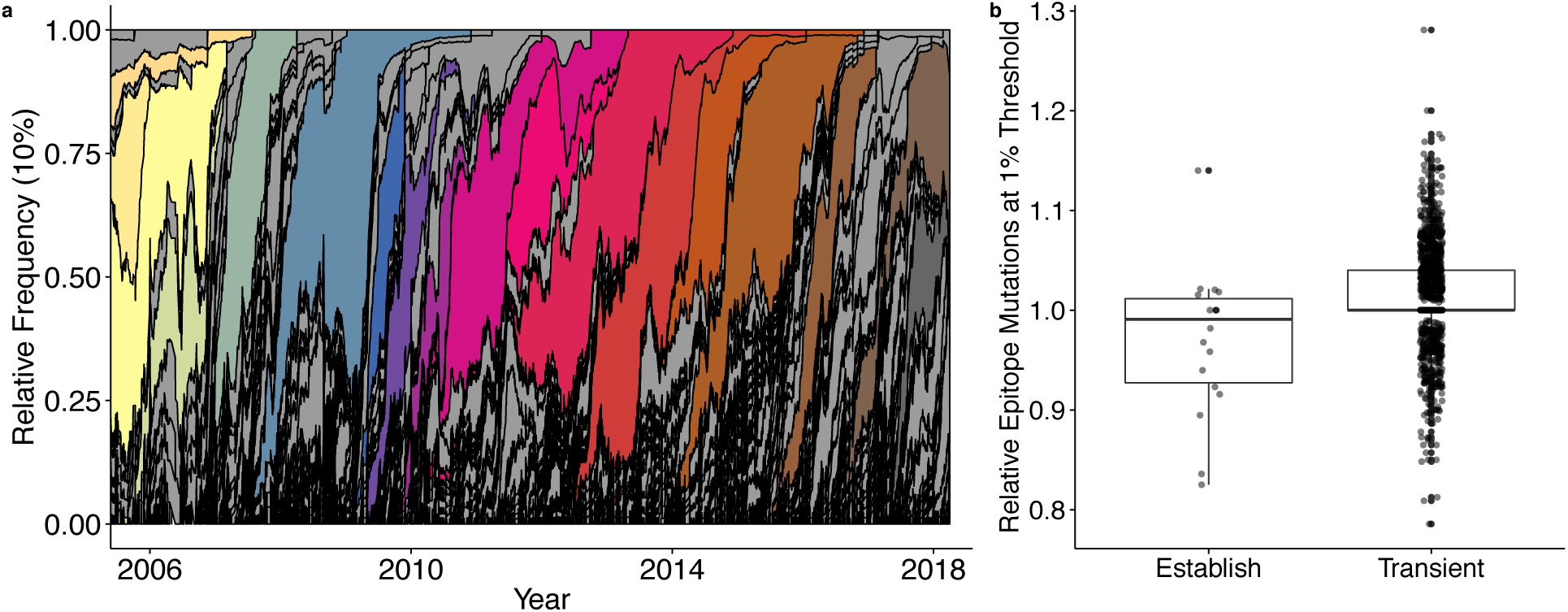
Empirical antigenic dynamics of influenza A/H3N2, 2006-2018. (a) Relative frequencies of all antigenic clusters that reach the threshold of at least 10% of sampled viruses. Frequencies are calculated using a 60-day sliding window. Grey shading indicates clusters that surpass the 10% threshold, but do not eventually establish (i.e., reach relative frequency of at least 20% for at least 45 days). Other colors indicate distinct antigenic clusters that eventually establish. (b) A low relative number of epitope mutations when a cluster reaches the 1% relative frequency threshold is an early indicator of future success (Wilcox, p < 0.003). We divide the number of epitope mutations of a focal cluster by the average number of mutations of simultaneously circulating clusters.

When forecasting influenza dynamics, there may be tradeoffs between prediction certainty, the extent advanced warning, and the surveillance effort required to detect and characterize emerging viruses. Across our ten models, there is a marked trade-off between lead-time and reliability, with low surveillance thresholds providing earlier but less accurate indicatication of future threats (Fig. 5). Across simulations, the median time difference between a cluster reaching the 1% and 10% surveillance thresholds was approximately 7 months (IQR: 154-294 days).

**Figure 5:**
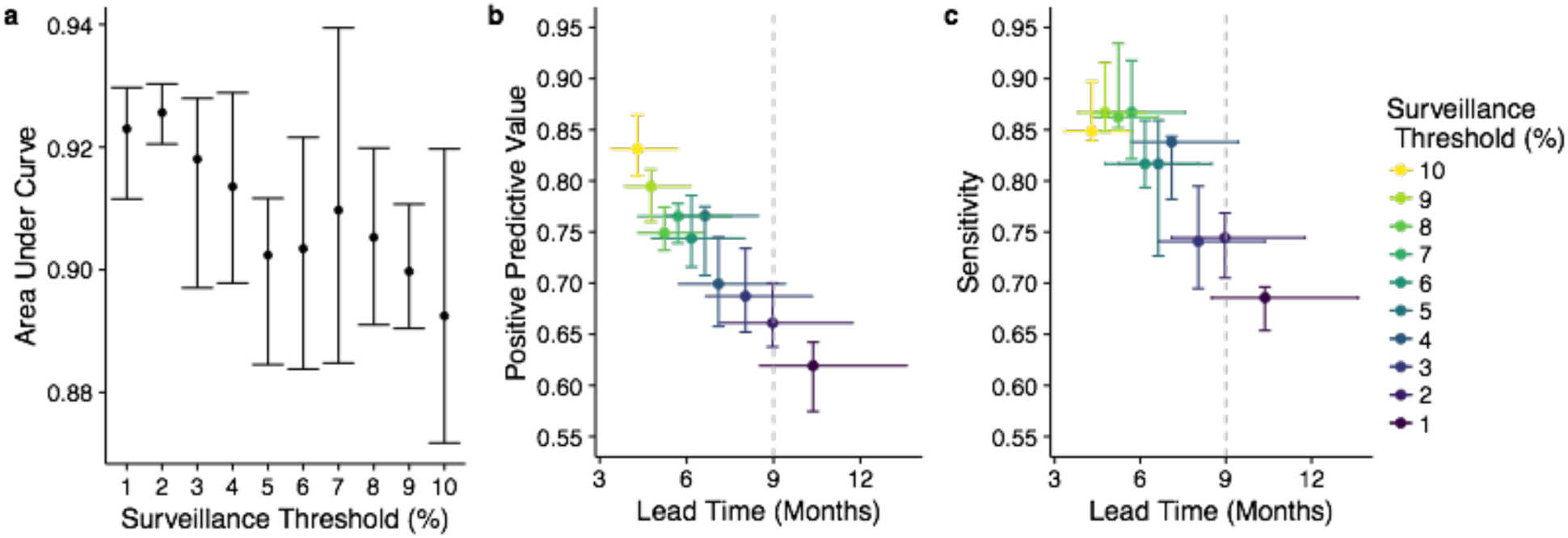
Model performance across surveillance thresholds. (a) Area under the receiver operator curve (AUC) suggests that models can predict successful from unsuccessful clusters by the time they reach 1% of circulating viruses, with discriminatory power declining slightly as clusters rise in frequency. Bars represent the max, median, and minimum AUC values across 5-fold cross validation. (b-c) There is a trade-off between lead time and model performance. The horizontal bars represent the IQR of time between the moment the expanding antigenic cluster reaches the surveillance threshold and when it reaches the success criteria. Vertical bars represent the range and median positive predictive value (b) and sensitivity (c), across five-fold evaluation. Colors correspond to the best fit model for each surveillance threshold. Dashed gray lines indicate lead times of nine months, which represents the current time between the Northern Hempisphere vaccine composition meeting in February and the following start of the influenza season in October.

Classifier models have substantial discriminatory and predictive power even when an antigenic cluster is present at low frequencies (Fig. 5a). Model AUC’s tend to decrease as the frequency of the candidate clusters increases. Conversely, the positive predictive value (PPV) and sensitivity increase at higher surveillance thresholds. The gains in sensitivity and PPV per month decrease at higher surveillance thresholds. Between the 1% and 5% surveillance thresholds, there is on average a 4% increase in sensitivity and 4.5% increase in positive predictive value per month lost in lead-time. However, between the 6% and 10% surveillance thresholds, sensitivity gains drop to 1.2% and ppv to 3.6% per month lost in lead-time. This decreasing tradeoff between gain in certainty and loss of lead-time reflects shorter intervals between surveillance thresholds as the cluster begins to rapidly expand and the model’s prediction capabilities reach upper capacity.

The primary predictor across all models––the relative growth rate of a cluster––cannot easily be estimated from available surveillance data. Thus, we built and evaluated bivariate logistic regression models that predict future success using more easily attained proxies (Table 2). One considers the time taken for the cluster to rise from 6% to 10% relative frequency and the total number of clusters that grew during this period; the other considers the fold-change in the relative frequency of the cluster between these time points and the background variance in fold-change. Of the four proxies, all but the relative fold-change of the cluster were statistically significant predictors, with negative effects on the probability of cluster success (Fig. S4). These resulting models have higher sensitivity than positive predictive values. We also tested analogous models using statistics calculated at alternative surveillance checkpoints (1% to 5%, 3% to 5%, and 8% to 10%), and found that the 6%-10% comparison performed best (Table S3).

**Table 2:** Performance of proxy predictors at the 10% surveillance threshold. Model 1 predicts the fate of a cluster using the top two predictors in our best fit model. The two proxy models use data from two time points, when the cluster reached relative frequencies of 6% (*t*_1_) and 10% (*t*_2_). Model 2 considers the time elapsed between and the number of competing expanding clusters. Model 3 considers the relative fold change in the focal cluster between the two time points and the population-wide variance in fold change. Performance values are the median of five fold cross-validation.

| Model | Type | AUC | PPV | Sensitivity |
| --- | --- | --- | --- | --- |
| 1. $R_c/\langle R \rangle + \text{var}(R)$ | Actual | 0.88 | 0.81 | 0.89 |
| 2. $\delta_c(t_1, t_2) + N_{\Delta_j(t_1, t_2) > 1}$ | Proxy | 0.78 | 0.74 | 0.87 |
| 3. $\chi_c(t_1, t_2) + \text{var}(\Delta_j(t_1, t_2))$ | Proxy | 0.67 | 0.66 | 0.95 |

Finally, we fit models to predict the presence and frequency of clusters based on opportunistic sampling of clusters, rather than waiting for specified surveillance thresholds. Cluster frequencies tend to skew towards low frequencies (Fig. S5). Our best fit model for predicting the future success of all clusters present at a random time point performs comparable to our best models for low surveillance thresholds (Fig. S6). We fit a second two-part model that sequentially predicts the presence-absence and the frequency of a cluster in three month intervals out to one year ahead. The model predicted up to twelve-month ahead presence-absence with 92% discriminatory power (AUC). However, the accuracy of the frequency predictions declined after six months, with a tendency to underestimate the frequencies of future dominant clusters (Fig. S7, Table S4-S5). The top predictors included the frequency of the cluster at the time of sampling and most of the top predictors selected for the surveillance threshold models.

## DISCUSSION

Until we develop an effective universal flu vaccine, seasonal vaccines will remain the frontline of influenza prevention. The severe 2017-2018 influenza season was a stark reminder that anticipating dominant strains with sufficient lead time for incorporation into vaccines is paramount to public health. Here, we analyzed over 1500 years of simulated influenza phylodynamics to explore the predictability of antigenic emergence and identify early predictors of future evolutionary success that can be plausibly monitored via ongoing surveillance efforts.

Phylodynamic models provide insight into both the interplay of evolutionary and epidemiological processes and how these dynamics are manifested in observable data. Our simulations revealed a stereotypical path to antigenic turnover consistent with those described in Koelle & Rasmussen [31]. An antigenic mutation appears on a virus. If its fitness is high relative to the competition, it can gain a foothold. In general, the lower the deleterious load and higher the susceptibility in the host population, the higher the fitness of the new virus. Thus, antigenic mutations occurring on good genetic backgrounds are more likely to gain traction [31]. The dynamics of susceptibility are a bit more complex. Although all hosts will be partially susceptible to a new antigenic type, the level of susceptibility will depend on past infections by antigenically similar viruses. We find that some successful mutants arise with markedly higher fitness than other co-circulating viruses, which propels them towards dominance, while others enter with only moderately high fitness but are able to ascend in the wake of a prior antigenic sweep, which suppresses other potentially competing viruses. Many of the mutants that eventually establish as dominant clusters first appear as another dominant cluster is cresting [44,45]. The rampant transmission affords opportunities for such mutations to arise and quashes other potentially competing clusters. Transmission of the previous cluster type begins to decreases as the population gains immunity through infection. With fewer susceptible hosts available to the previously dominant cluster, the expanding cluster, as well as any competing clusters, begin to constitute a larger fraction of all circulating viruses.

As a cluster expands, it accumulates deleterious mutations. By the time new clusters reach a relative frequency of 10%, their mutational loads begins to approach the population average. Simultaneously, the number of susceptible hosts decreases as the cluster sweeps through the host population. If a cluster reaches a relative frequency of 10%, its probability of future dominance will be influenced by how much of a fitness advantage it retains, and by the level of competition from other clusters.

The strongest predictor of future dominance across all of our models is the relative effective reproductive number of a cluster, that is, the growth rate of the cluster compared to the average growth rate across the viral population. This measure of viral fitness incorporates both the real-time competitive advantage (vis-a-vis the immunological landscape) and deleterious mutational load. Intuitively, faster growing clusters are more likely to persist and expand. Our ability to predict the fate of an emergent virus improves as the cluster increases in relative frequency. Both sensitivity––the proportion of successful clusters detected by the model––and positive predictive value––the proportion of predicted successes that actually establish––surpass 80% by the time a cluster has reached 10% relative frequency.

The second most informative predictor selected across all models––the population-wide variance in the effective growth rate, var(*R*)––requires a more nuanced interpretation. The greater the background variance at the time a cluster is emerging, the less likely the cluster is to succeed. To unpack this result, we analyzed the competitive environment of emerging clusters with only modest growth rates; rapidly growing clusters are likely to succeed regardless of their competition. Within this class of slowly emerging viruses, those that initially face a single high frequency dominant cluster and fewer co-emerging competitors are more likely to succeed [46,47]. A recent sweep by a dominant cluster leaves a wake of immunity that can be exploited by antigenically-novel clusters that stochastically battle for future dominance. We hypothesize that these two conditions––a reigning dominant cluster and reduced competition with emerging novelty––reduce the overall variance in viral growth rate and explain the negative correlation between this quantity and the future ascent of an emerging cluster.

While the certainty of our predictions improves as clusters increase in relative frequency, there is a trade-off with lead time. The longer we wait to assess a rising cluster, the less time there will be to update vaccines and implement other intervention measures. For a successful cluster detected at a relative frequency of 1%, there will be, on average, 10 months before the cluster becomes established (maintains a relative frequency over 20% for 45 days). If detected only after reaching a relative frequency of 10%, the expected lead time shrinks to four months. Although real-world surveillance is noisy and dependent on sufficient sampling depth and geographic coverage, our results suggest that, with a perfect knowledge of the host and viral populations, predictions can be made with at least 85% sensitivity and confidence before a cluster rises to 10% of all circulating strains.

As policy-makers consider new strategies for antigenic surveillance and forecasting, the trade-off between prediction accuracy and lead time has practical implications. For example, a detection system targeting new viruses as soon as they reach 1% relative frequency has the benefit of early warning and drawback of low accuracy, which translate into economic and humanitarian costs and benefits. On the positive side, early warning increases the probability that seasonal vaccines will provide a good match with circulating strains, and thus lowers the expected future morbidity and mortality attributable to seasonal influenza. Based on the vaccine production and delivery schedule, the surveillance window for emerging clades is from October to February for the Northern Hemisphere and vaccine composition is determined at an international meeting in February [17]. Our analysis suggests that, at nine months before an emerging cluster sweeps to dominance, it is likely to have be circulating at a low relative frequency in the range of 1% to 4%. On the negative side, the low surveillance threshold for candidate clusters and consequent lower accuracy require far more surveillance and vaccine development resources than higher surveillance thresholds. In our simulations, for example, the number of clusters screened at the 1% threshold is an order of magnitude higher than at the 10% surveillance threshold and the number of false positive predictions potentially prompting further investigation is also manifold greater.

Our top predictors of viral emergence require a comprehensive sampling of the viral and host population. Although exact measurements of these quantities are practically infeasible, our results suggest that targeting molecular surveillance towards precise and accurate estimation of viral growth rates, both for newly emerging clusters and the resident circulating viruses, may enhance influenza prediction. One approach is to target the two key components of growth rate separately––mutational load and effective susceptibility. Changes in the mutational load can be estimated from sequence data, comparing the number of differences that occur in non-epitope portions of the genome over time [23,27,48]. Our parameterization of *R*_*c*_ follows the empirical method of [23], with fitness costs based on nonsynonymous amino acid differences between a given strain and its most recent common ancestor. Estimating the effective susceptibility is more challenging, as it depends on the interaction between an individual’s exposure history [49–51] and new amino acid substituions in epitode coding regions [6,52]. Nonetheless, several studies introduce innovative methods for estimating susceptibility from the historic distribution of influenza subtypes, seasonal influenza prevalence, and HI-titers. For example, Neher *et al*. [48] predict antigenic properties of novel clades by mapping both serological and sequence data to a phylogenetic tree structure of HA sequences. Łuksza & Lässig estimate effective susceptibility by first estimating the historic frequency of clades in six-month intervals and then estimating cross-immunity between those clades and the focal cluster based on amino acid differences in epitope regions [23]. However, both methods only consider clusters that have already surpassed 10% relative frequency, at which point strains are thought to be geographically well-mixed and less prone to geographic sampling bias.

Another approach to estimating the growth rate of an emerging cluster is to treat it as a composite quantity. We evaluated several proxy measures of cluster growth rate, including the relative fold-change in frequency between two time points. Models based on fold change rather than the true growth rate actually have greater sensitivity, that is, they are more likely to detect clusters destined for dominance when they first emerge. However, the positive predictive values of our best models drop from from 0.81 to 0.67, meaning that replacing the true growth rates with an approximation increases the rate of false alarms. Importantly, the proxy model improves with the addition of a second predictor, the variance in fold-change across the viral population, which can also be readily estimated from surveillance data. Thus, variance in fitness appears to be a robust secondary predictor of future sweeps, regardless of how fitness is quantified. Surprisingly, a model based on seemingly naive approximations of growth rate––the time elapsed between two frequency thresholds and the number of other co-circulating clusters rising in frequency––was even more accurate, though still inferior to the true growth rate models. We did not evaluate a promising alternative strategy for approximating fitness, based on the phylogenetic reconstruction of currently circulating sequences [25,53]. Unlike the proxies we considered, this does not require historical data but does rely on pathogen sequencing. Finally, although not as informative as predictors that quantify the evolutionary and immunological state of the population, easily quantifiable predictors such as the total number of infected individuals or the frequency of the circulating dominant cluster, can be incorporated into future predictive models.

Our attempts to directly apply the optimized models to empirical data were of limited success. While the global evolutionary dynamics of influenza A/H3N2 clusters visually resemble those observed in our simulations, the sparse genotypic data available do not permit estimation of the phenotypic predictors identified in our study. The number of epitope mutations in a newly emerging cluster relative to co-circulating viruses provides early indication of future success. This provides proof of concept that the evolutionary viability of influenza viruses is predictable, but will require better models for estimating viral fitness from sequence data and the expansion of surveillance efforts [54] to collect phenotypic data reflecting the mutational loads viruses and dynamic trends in population susceptibility.

While our study provides actionable suggestions for improving both the surveillance and forecasting of antigenic turnover, it is limited by several assumptions. One caveat of our method is that we do not capture the explicit phylogenetic structure of the influenza population. Therefore, we do not distinguish between clusters that are successful because of one mutation and clusters that are successful because of a series of mutations. If for instance, a novel antigenic mutation caused the emergence of a new cluster (phenotype) that circulated briefly before a second novel antigenic mutation caused a second phenotype that eventually achieved our defined criteria, we ignore the fact that the established cluster is a subclade of the first and that the antigenic mutation that conferred the first phenotype is fixed along with the second antigenic mutation [46,55]. This scenario follows Koelle & Rasmussen’s description of a two-step antigenic change molecular pathway that leads to antigenic cluster transitions [31]. Our analysis is therefore relevant for scenarios that depict their described jackpot strategy -- a combination of one large antigenic mutation occuring on a low deleterious background. Second, the simulation represents global H3N2 dynamics and ignores differences in temperate and tropical transmission dynamics [34,45,56]. Prior studies have revealed considerable global variation in transmission rates, which should positively correlate with the frequency of cluster transitions. Furthermore, viruses that emerge in tropical regions are more likely to be the source of viruses that eventually circulate in temperate regions [57,58]. Temperate regions produce more extreme seasonal bottlenecks, potentially leading to greater stochasticity in viral dynamics, which makes it more difficult for novel strains that emerge in temperate regions to spread globally [59]. We also do not consider selective pressures imposed by seasonal vaccination. Its impact on antigenic turnover depends on vaccination rates and the immunological match between the vaccine and all co-circulating viruses. Seasonal vaccination could differentially modify the effective susceptibility of clusters, suppressing some while creating competitive vacuums for others. Theoretical study suggests that antigenic drift should slow down [60,61] and the circulation of co-dominant clusters may become more common [62]. Given these caveats, we emphasize our qualitative rather than the quantitative results. Our study highlights promising predictors of viral success, characterizes robust trade-offs between the timing, costs and accuracy of such predictions, and serves as proof-of-concept that model-derived surveillance strategies can accelerate and improve forecasts of antigenic sweeps. If we fit similar models directly to historic surveillance data, the resulting predictions will likely reflect greater uncertainty but perhaps naturally reflect global variation in influenza dynamics and vaccination pressures.

Our study demonstrates that the early detection of emerging influenza viruses is limited by a tight race between the typical dynamics of antigenic turnover and the annual timeline for influenza vaccine development. It also provides a foundation for analyzing the costs and benefits of expanding surveillance capacities and shortening the vaccine production pipeline. As we strive to expedite and improve molecular surveillance for vaccine strain selection, even incremental progress is valuable. Earlier detection of antigenic sweeps, regardless of vaccine efficacy, can inform better predictions of severity, public health messaging regarding personal protective measures, and clinical preparedness for seasonal influenza.

## Supporting information

Data Acknowledgments

## Acknowledgements

We are grateful to John Huddleston for providing the empirical data for the phylogenetic analysis. We acknolwedge the authors, originating and submitting laboratories of the sequences from GISAID’s EpiFlu Database (see attached table for sequence accession details). We acknowledge the Texas Advanced Computing Center (TACC) at The University of Texas at Austin for providing High performance computing resources that have contributed to the research results reported within this paper. URL: http://www.tacc.utexas.edu. LAC was supported by the Department of Defense (DoD) through the National Defense Science & Engineering Graduate Fellowship (NDSEG) Program. TB was supported by NIH NIGMS R35 GM119774-01 NIH NIAID U19 AI117891. TB is a Pew Biomedical Scholar. LAM was supported by the National Institutes of Health (Models of Infectious Disease Agent Study grant U01 GM087719).

## Author Contributions

LAC and LAM developed the conceptual framework and study design. LAC, TB, and LAM conceived of the analyses. LAC performed the analyses, created the figures, and wrote the first draft. LAC, LAM, and TB interpreted all results. All authors contributed to the writing and approval of the final manuscript.

## Data and Code Availability

Source code for the full phylodynamic model that produced the data in this study is available on GitHub at https://github.com/davidrasm/MutAntiGen.git. The phylodynamic model in this paper is a modified version of the program Antigen (http://bedford.io/projects/antigen/). The list of sequences used in the empirical analysis is included in the acknowledgements table.

