## Supplemental Material for "Early Prediction of Antigenic Transitions for Influenza A H3N2"

### Supplement: Early Prediction of Antigenic Transitions for Influenza A H3N2: Electronic supplementary material

#### 1 Extended Methods

##### 1.1 Influenza Phylodynamic Simulations

**Deriving the criteria for cluster establishment in the population**

To separate the clusters that are eventually successful from those that only transiently circulate, we derived a two-criteria threshold of establishment based on reaching a minimum frequency in the population, and circulating above the minimum frequency for a specified duration of time. We choose the most stringent criteria that, when only accounting for clusters that reached the criteria, still maintained cyclical influenza dynamics. In addition, we compared the proportion of the total infected population attributable to established clusters under different criteria (Fig. 1).

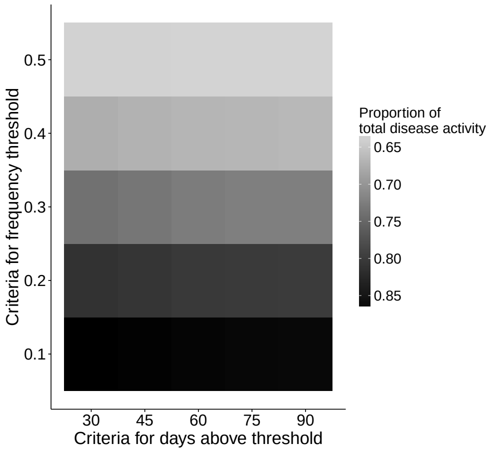


Figure 1: **The proportion of total infections caused by established clusters is more sensitive to a frequency criteria than the duration of time**. Established antigenic clusters account for the majority of the disease activity at any point in time. In our analysis, established clusters are those that circulate above 20% relative frequency for at least 45 days. Using this criteria, infections caused by future successful clusters account for a median of 81% of the disease activity at any point in time.

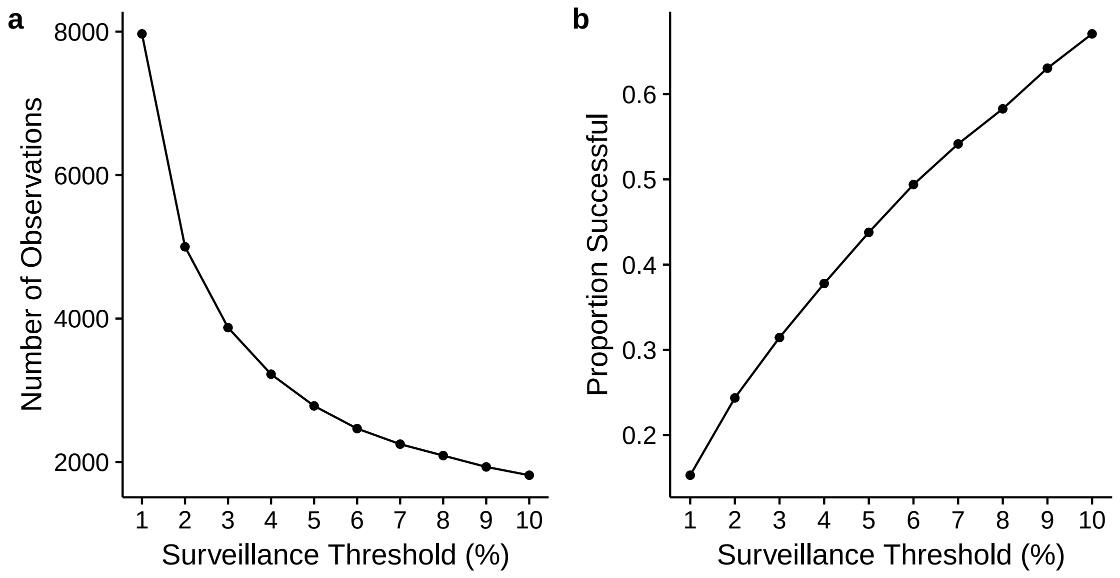


Figure 2: **The fate of novel antigenic clusters** (A) Each point represents the number of antigenic clusters in our simulations that reach each increasing surveillance threshold (i.e., relative frequency in the population). As the surveillance threshold increases from 1 to 10%, the number of candidate clusters decreases from 7969 clusters at the 1% threshold to 1816 clusters at the 10% surveillance threshold. (B) Given a cluster has reached a surveillance threshold, the proportion of antigenic clusters that will establish (i.e. reaches $> 20\%$ for 45 days) increases with higher surveillance thresholds.

#### 1.2 Candidate Predictors

Candidate predictors are those population genetic and epidemiological indicators that were tracked during the phylodynamic simulations or calculated from the output. The full list is in Table 1.

| Candidate Predictor | Formula | Population | Cluster | Relative |
|---|---|---|---|---|
| Number of Infected Individuals | $I$ | X | | |
| Number of Uninfected Individuals | $S$ | X | | |
| Proportion of Individuals Infected | $I/N$ | X | | |
| Number of Circulating Antigenic Clusters | $N_c$ | X | | |
| Frequency of Current Dominant Cluster | $f_c = \max\left[I_c/I\right]$ | X | | |
| Entropy (Shannon's Diversity Index) | $H = \frac{1}{N_C}\sum_{j=1}^{N_C} f_j \ln \frac{1}{f_j}$ | X | | |
| Serial Interval of Infection* | $\mathrm{SI} = \frac{1}{I}\sum_{a,b\in\text{infecteds}} \left(t_{a_0} - t_{b_0}\right)$ | X | | |
| The most recent common ancestor* | $\mathrm{TMRCA} = \max\left[\frac{1}{2}\left(\left(t_{TMRCA_0} - t_{a_0}\right) + \left(t_{TMRCA_0} - t_{b_0}\right)\right)\right]$ | X | | |
| Genetic Diversity* | $\omega = \frac{1}{I}\sum_{a,b\in\text{infecteds}} \frac{1}{2}\left(\left(t_{TMRCA_0} - t_{a_0}\right) + \left(t_{TMRCA_0} - t_{b_0}\right)\right)$ | X | | |
| Antigenic Diversity* | $\lambda = \frac{1}{I}\sum_{a,b\in\text{infecteds}} \lambda_{ab}$ | X | | |
| Deleterious Mutational Load | $\mathrm{k} = \frac{1}{I}\sum_{i=1}^{I} k(v_i)$ | Mean, Var | Mean, Var | Mean, Var |
| Transmissibility | $\beta = \frac{1}{I}\sum_{i=1}^{I} \beta_0\left(1 - s_d\right)^{k(v_i)}$ | Mean, Var | Mean, Var | Mean, Var |
| Effective Susceptibility* | $S_{\text{eff}}(v) = \frac{S}{N}\sum_{h=1}^{N}\left(\sigma_{v(h)}\right)$ | Mean, Var | | |
| Covariance in transmissibility and effective susceptibility | $\mathrm{cov} = \frac{1}{I}\sum_{i=1}^{I}\left(\left(\beta_i - \bar{\beta}\right) * \left(\sigma_v(i) - \bar{\sigma}\right)\right)$ | X | | |
| Cluster Susceptibility* | $\sigma(v) = \sum_{h=1}^{N}\min(1, \sigma_{v,c(h,v)})$ | | Mean, Var | |
| Reproductive Growth Rate | $R(v) = \frac{\beta_0\left(1-s_d\right)^{k(v)}}{\mu+\nu}\left(\frac{S_{\text{eff}}(v)}{N}\right)$ | Mean, Var | Mean, Var | Mean, Var |

Table 1: **Full set of candidate predictors considered**. Values were taken at the moment a focal antigenic cluster reached a specified surveillance threshold. The columns *Population, Cluster, Relative* indicate the scale and measure (e.g. mean and/or variance) that a predictor was considered in the model. Depending on the scale of the predictor, the *formula* could refer to all strains in the population, i.e. the strains of infected hosts, or the subset of strains in a specific cluster. *For computational simplicity, these quantities were calculated using strains from a random sample of 10,000 infected individuals. N = number of hosts; $t_{a_0}$ = the time of birth of virus $a$; $\lambda$ = antigenic distance between two strains. The antigenic distance is the pairwise degree of cross-immunity between two strains determined by the size of antigenic mutations and parent-offspring relationships; $k(v_i)$ = the number of deleterious mutations on a virus $v$ of infected host $i$; $s_d$ = the fitness effect of a deleterious mutation; $\sigma_v$ = the average individual population susceptibility to cluster $c$; $\sigma_{v,c(h,v)}$ = the probability of infection of a host with historical infection $i$ by a strain of cluster $v$

### 2 Results

#### 2.1 Incorporation of historical data

We tested an additional strategy for building classifier models: one that incorporated previously sampled data for each cluster.

To predict a cluster's evolutionary outcome for a specific surveillance threshold, we used data from three timepoints: 1) when it reached 1%, 2) when it reached the half-way frequency level between 1% and the surveillance threshold of interest, and 3) when it reached the surveillance threshold. The candidate predictors included all variables listed in Table 1, as well as the difference in predictor values between 1% and the halfway point, and the halfway point and the maximum surveillance threshold. In addition, the difference of these differences was an additional predictor. Models were fit using the same approach described in the manuscript.

To compare the performance of models that incorporated data from past time points to models based only on current data, we compared the sensitivity and positive predictive value across surveillance thresholds from $1 - 10\%$. Both model types performed similarly across the range of surveillance thresholds (Fig. 3). we focus on strategy that only incorporates current data because of the simplicity in the methodology and reduction of candidate predictors.

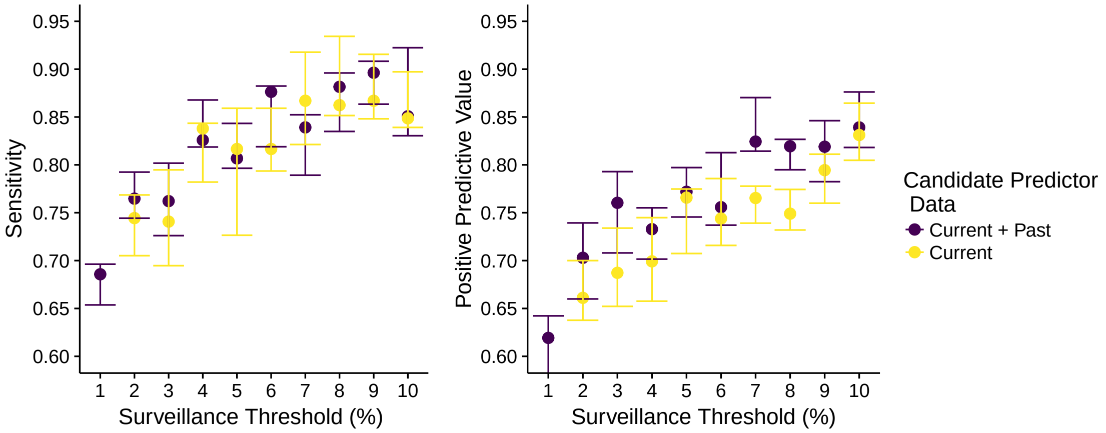


Figure 3: **Predictive models that rely on data from a single sample of time perform similarly to those that include data from multiple time points**. Each point represents the combination of candidate predictors that best predict the evolutionary fate of antigenic clusters at varying surveillance thresholds, whether the combination includes data from a single time point (yellow) or multiple time points (purple). Model performance is measured in terms of sensitivity and positive predictive value. Dots represent the median, and error bars span the range of performance values across the five folds of cross-validation of the best-fit model.

#### 2.2 Surveillance Threshold Results

The best-fit logistic regression models for surveillance thresholds from $1 - 10\%$ are listed in Table 2.

| Surveillance Threshold (%) | Predictor Variable | Coefficient Estimate | Std. Error |
|---|---|---|---|
| 1 | $R_c/\langle R\rangle$ | [2.61, 2.77] | [0.09,0.09] |
| | var($R$) | [-0.54, -0.60)] | [0.06,0.06] |
| | $\langle R\rangle$ | [0.30, 0.40] | [0.05-0.05] |
| | $k_c/\langle k\rangle$ | [-0.20, -0.28)] | [0.06, 0.06] |
| | var($\sigma_c$)/var($S_{\text{eff}}$) | [0.12, 0.16] | [0.05-0.06] |
| | var($\beta_c$)/var($\beta$) | [0.17, 0.24] | [0.05-0.05] |
| | var($\sigma_c$) | [0.11, 0.19] | [0.05-0.06] |
| 2 | $R_c/\langle R\rangle$ | [2.60, 2.71] | [0.09-0.10] |
| | var($R$) | [-0.52, -0.57] | [0.06-0.07] |
| | $\langle R\rangle$ | [0.28, 0.40] | [0.05-0.05] |
| | $k_c/\langle k\rangle$ | [-0.27, -0.34] | [0.06-0.06] |
| | var($\beta_c$)/var($\beta$) | [0.32, 0.36] | [0.05-0.06] |
| | I/N | [-0.16, 0.18] | [0.06-0.06] |
| | var($\sigma_c$) | [0.13, 0.19] | [0.06-0.06] |
| 3 | $R_c/\langle R\rangle$ | [2.36, 2.43] | [0.09-0.10] |
| | var($R$) | [-0.46, -0.57] | [0.07-0.07] |
| | $\langle R\rangle$ | [0.39, 0.47] | [0.06-0.06] |
| | $k_c/\langle k\rangle$ | [-0.32, -0.37] | [0.06-0.06] |
| | var($\beta_c$)/var($\beta$) | [0.26, 0.29] | [0.06-0.06] |
| | I | [-0.14, 0.21] | [0.06-0.06] |
| | max$[[I_c//I_t]$ | [0.11, 0.19] | [0.06-0.06] |
| 4 | $R_c/\langle R\rangle$ | [2.53-2.75] | [0.10-0.11] |
| | var($R$) | [-0.47, -0.63] | [0.07-0.08] |
| | $\langle R\rangle$ | [0.30, 0.42] | [0.06-0.06] |
| | $k_c/\langle k\rangle$ | [-0.22, -0.28] | [0.06-0.07] |
| | var($\beta_c$)/var($\beta$) | [0.30, 0.33] | [0.06-0.06] |
| | var($\sigma_c$) | [0.15, 0.22] | [0.06-0.06] |
| | I | [-0.15, 0.17] | [0.06-0.06] |
| 5 | $R_c/\langle R\rangle$ | [2.35, 2.58] | [0.11-0.12] |
| | var($R$) | [-0.46, -0.59] | [0.08-0.08] |
| | $\langle R\rangle$ | [0.31, 0.44] | [0.06-0.06] |
| | $k_c/\langle k\rangle$ | [-0.25, -0.32] | [0.06-0.07] |
| | var($\sigma_c$) | [0.17, 0.23] | [0.06-0.06] |
| | var($\beta_c$)/var($\beta$) | [0.14, 0.20] | [0.06-0.06] |
| | max$[I_c/I_t]$ | [0.11, 0.17] | [0.06-0.06] |
| 6 | $R_c/\langle R\rangle$ | [2.36, 2.60] | [0.11-0.12] |
| | var($R$) | [-0.60, -0.71] | [0.08-0.08] |
| | $\langle R\rangle$ | [0.32, 0.43] | [0.06-0.06] |
| | $k_c/\langle k\rangle$ | [-0.26, -0.17] | [0.06-0.06] |
| | var($\beta_c$)/var($\beta$) | [0.19, 0.23] | [0.06-0.06] |
| | var($\sigma_c$) | [0.09, 0.21] | [0.06-0.07] |
| 7 | $R_c/\langle R\rangle$ | [2.36, 2.59] | [0.12-0.13] |
| | var($R$) | [-0.69, -0.78] | [0.08-0.08] |
| | $\langle R\rangle$ | [0.32, 0.40] | [0.07-0.07] |
| | $k_c/\langle k\rangle$ | [-0.25, -0.32] | [0.07-0.07] |
| | var($\beta_c$)/var($\beta$) | [0.17, 0.25] | [0.06-0.07] |
| 8 | $R_c/\langle R\rangle$ | [2.22, 2.40] | [0.11-0.12] |
| | var($R$) | [-0.51, -0.62] | [0.08-0.09] |
| | $\langle R\rangle$ | [0.32, 0.42] | [0.07-0.07] |
| | $k_c/\langle k\rangle$ | [-0.27, 0.34] | [0.07-0.07] |
| | var($\beta_c$)/var($\beta$) | [0.17, 0.25] | [0.07-0.07] |
| | max$[I_c/I_t]$ | [0.15, 0.29] | [0.07-0.07] |
| 9 | $R_c/\langle R\rangle$ | [2.24, 2.45] | [0.12-0.13] |
| | var($R$) | [-0.48, -0.63] | [0.09-0.10] |
| | $\langle R\rangle$ | [0.30, 0.38] | [0.07.-0.07] |
| | $k_c/\langle k\rangle$ | [-0.22, 0.31] | [0.07-0.07] |
| | max$[I_c/I_t]$ | [0.13, 0.25] | [0.07-0.07] |
| | var($\beta_c$)/var($\beta$) | [0.09, 0.24] | [0.07-0.07] |
| 10 | $R_c/\langle R\rangle$ | [2.38, 2.55] | [0.13-0.14] |
| | var($R$) | [-0.63, -0.74] | [0.09-0.09] |
| | $\langle R\rangle$ | [0.27, 0.35] | [0.07-0.07] |
| | $k_c/\langle k\rangle$ | [-0.18, -0.23] | [0.07-0.08] |
| | var($\beta_c$)/var($\beta$) | [0.14, 0.22] | [0.08-0.08] |
| | TMRCA | [-0.10, -0.21] | [0.08-0.08] |

Table 2: **Best-fit model results for surveillance thresholds** $1-10\%$**.** The predictor variables are listed in the order by which they were selected using a forward selection algorithm. The coefficient estimate is the maximum and minimum coefficient (log-odds) from the five-fold cross validation of the final full-term model with the corresponding std. error.

#### 2.3 Proxy Model Results

Because the top selected predictors across all models cannot easily be estimated using readily available surveillance data, we evaluated several proxy measures of viral growth rates and viral competition. The results of the models and how they compare to the performance of a model using only the true relative epidemiological growth rate and the background variance in growth rate can be seen in Table 3. In addition to the terms included in the table, we tested the fold change of the dominant cluster from $t_1$ and $t_2$as a predictor, but did not find that this term was a significant proxy in any model.

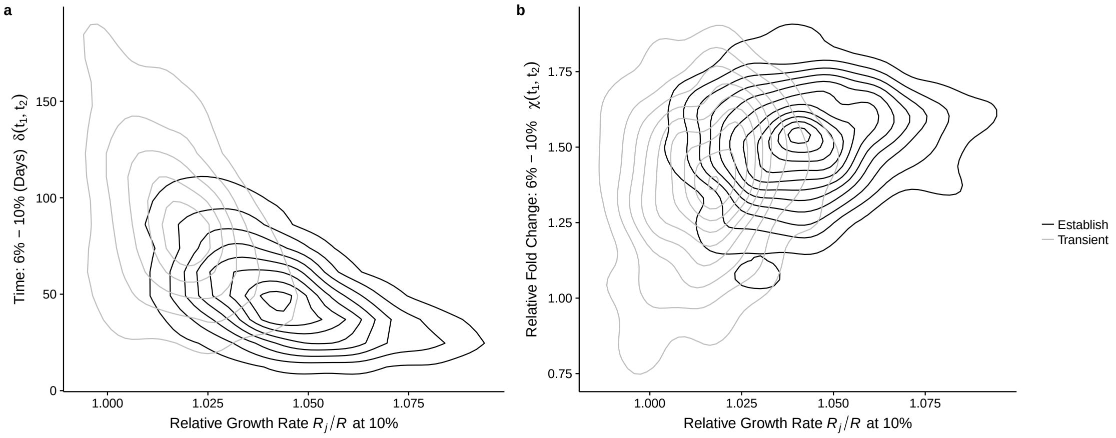


Figure 4: **The rate of change (a) and relative fold change (b) as proxy measures for the relative growth rate** $R/\langle R\rangle$. Contour lines indicate the density of cluster values for clusters that establish (black) and those that transiently circulate (grey). Values along the x-axis indicate the measured relative growth rate of a cluster the moment it reaches the 10% surveillance threshold. Values along the y-axis indicate the proxy measure (rate of change in (a) and relative fold change in (b) for the cluster, approximated for the time between the 6% and 10% thresholds. The rate of change, measured in the number of days between the two thresholds, is a better proxy measure than relative fold change .

| **Surveillance Thresholds** | **Model** | Type | **Balanced Accuracy** | **AUC** | **PPV** | **Sensitivity** |
|---|---|---|---|---|---|---|
| 5% | 1. $R_c/\langle R\rangle$ + var($R$) | Actual | 0.78 | 0.88 | 0.81 | 0.89 |
| 1-5% | 2. $\delta_c(t_1, t_2)$ + $N_{\Delta_j(t_1,t_2)>1}$ | Proxy | 0.57 | 0.71 | 0.65 | 0.93 |
| 1-5% | 3. $\chi_c(t_1, t_2)$ + var($\Delta_j(t_1, t_2)$) | Proxy | 0.50 | 0.58 | 0.61 | 0.99 |
| 5% | 1. $R_c/\langle R\rangle$ + var($R$) | Actual | 0.78 | 0.88 | 0.81 | 0.89 |
| 3-5% | 2. $\delta_c(t_1, t_2)$ + $N_{\Delta_j(t_1,t_2)>1}$ | Proxy | 0.56 | 0.67 | 0.64 | 0.95 |
| 3-5% | 3. $\chi_c(t_1, t_2)$ + var($\Delta_j(t_1, t_2)$) | Proxy | 0.50 | 0.58 | 0.61 | 0.99 |
| 10% | 1.$R_c/\langle R\rangle$ + var($R$) | Actual | 0.78 | 0.88 | 0.81 | 0.89 |
| 6-10% | 2.$\delta_c(t_1, t_2)$ + $N_{\Delta_j(t_1,t_2)>1}$ | Proxy | 0.70 | 0.78 | 0.74 | 0.87 |
| 6-10% | 3.$\chi_c(t_1, t_2)$ + var($\Delta_j(t_1, t_2)$) | Proxy | 0.59 | 0.67 | 0.66 | 0.95 |
| 10% | 1. $R_c/\langle R\rangle$ + $\sigma_R$ | Actual | 0.78 | 0.88 | 0.81 | 0.89 |
| 8-10% | 2. $\delta_c(t_1, t_2)$ + $N_{\Delta_j(t_1,t_2)>1}$ | Proxy | 0.63 | 0.72 | 0.68 | 0.95 |
| 8-10% | 3. $\chi_c(t_1, t_2)$ + var($\Delta_j(t_1, t_2)$) | Proxy | 0.58 | 0.70 | 0.65 | 0.97 |

Table 3: **Evaluating proxy measures for different phases of a novel antigenic cluster's early expansion**. Model 1 shows the performance of the best-fit model using the actual values of relative fitness (relative growth rate) and competition (variance in the population growth rate) for clusters that reached the 5% surveillance thresholds (top two sections) and the 10% surveillance threshold (bottom two sections). Within each section, Model 2 substituted a time proxy for the fitness term and the absolute number of clusters that were growing for the competition term. Model 3 substituted a relative fold change for the fitness term and the population-wide variance in fold change for the competition term. $t_1$ is when a focal cluster reaches the lower surveillance threshold $(1\%, 3\%, 6\%, 8\%)$; $t_2$ is when the same cluster reaches the higher surveillance threshold $(5\%, 10\%)$ Performance metric values are the median across the five folds in cross-validation. Balanced accuracy measures the accuracy of the model, accounting for the imbalance in outcomes (i.e. number of transient versus established clusters) in the data set.

### 3 Alternate Surveillance Paradigms

Instead of a making antigenic transition predictions based on specific surveillance thresholds, one may opportunistically make predictions on cluster fate when samples become available. To compare the robustness of important predictors and model performance under this type of surveillance strategy, we fit two other model types, focused on 1) predicting the evolutionary fate of a cluster and 2) predicting the frequency up to a year out in 3-month increments.

Our data set consisted of all the antigenic clusters present in 10 random time points over a 25 year time period for each of the 62 independent simulations (N = 2846 clusters). We collected candidate predictors for all antigenic clusters that were present above 1% frequency and that had not already surpassed the successful criteria. In addition to the candidate predictors listed in Table1, the present frequency of each antigenic cluster, $f_c$, at the time of sampling was also included as a predictor. To directly compare the two types of surveillance strategies, we found the best fitting model that predicted the antigenic cluster's evolutionary fate using the same cross-validation model fitting process previously described 6.

Second, rather than a binary transient-successful classification, we predicted frequency levels of present transient clusters at 3 month intervals up to 12 months into the future. Out of the 2846 unique clusters in this data-set, 2279 cluster had $f_c' \geq 0.01\%$ at 3 months; 1921 clusters at 6 months, 1624 clusters at 9 months, and 1378 clusters at 12 months. For each 3 month increment, we went through the following model fitting process. First, classification models were built to assess whether an antigenic cluster would be present or absent in $X$ months time. Next, regression models were fit to predict the frequency for any antigenic cluster that was present above 1% in $X$ months time. At each step, candidate predictor values of the eligible clusters were centered and scaled. To improve model fit, the target frequency, $f_c'$ in $X$ months was log-transformed. We tested the performance of the best-fit model for each 3-month increment on a new data set consisting of 5 random time points over a 25 year period, corresponding to 310 time points over all 62 simulations (Tables 4, 5, Fig. 7). In addition, we tried fitting the model the frequency fold in X months time, i.e. $f_c(t+X)/f_c(t)$; however the model's goodness-of-fit, as measured by the adjusted $R^2$ was consistently lower than that of the models predicting the log-transformed frequency.

| Months Ahead | Predictors | AUC | PPV | Sensitivity |
|---|---|---|---|---|
| 3 | $f_c$<br>$R_c$<br>$\langle R \rangle$<br>$\beta_c/\langle\beta\rangle$<br>$\mathrm{var}(\beta_c)/\mathrm{var}(\beta)$ | 0.93 | 0.90 | 0.96 |
| 6 | $f_c$<br>$R_j$<br>$\langle R \rangle$<br>$\beta_c/\langle\beta\rangle$<br>$\mathrm{var}(R)$<br>$\mathrm{var}(\beta_c)/\mathrm{var}(\beta)$<br>$I$ | 0.93 | 0.89 | 0.89 |
| 9 | $R_c$<br>$f_c$<br>$\langle R \rangle$<br>$\beta_c/\langle\beta\rangle$<br>$\mathrm{var}(R)$<br>$\mathrm{var}(\beta_c)/\mathrm{var}(\beta)$<br>tMRCA | 0.93 | 0.84 | 0.91 |
| 12 | $R_j$<br>$f_c$<br>$\langle R \rangle$<br>$\mathrm{var}(R)$<br>$\beta_c/\langle\beta\rangle$<br>$\mathrm{var}(\beta_c)/\mathrm{var}(\beta)$ | 0.92 | 0.81 | 0.87 |

Table 4: Best-fit logistic regression results for predicting presence-absence of a cluster in X months time into the future. Terms are listed in the order they were added to the model through forward-selection.

| Months Ahead | Predictors | $R^2$Adjusted | RMSE (log) |
|---|---|---|---|
| 3 | $f_c$<br>$R_c$<br>$\langle R \rangle$<br>$\beta_c/\langle \beta \rangle$ | 0.84 | 0.36 |
| 6 | $f_c$<br>$R_c$<br>$\langle R \rangle$<br>$\beta_c/\langle \beta \rangle$<br>$\mathrm{var}(R)$ | 0.74 | 0.51 |
| 9 | $R_c$<br>$f_c$<br>$\langle R \rangle$<br>$\mathrm{var}(R)$<br>$\beta_c/\langle \beta \rangle$<br>$\mathrm{var}(\beta_c)/\mathrm{var}(\beta)$ | 0.66 | 0.65 |
| 12 | $R_j$<br>$f_c$<br>$\langle R \rangle$<br>$\mathrm{var}(R)$<br>$\mathrm{var}(\beta_c)/\mathrm{var}(\beta)$<br>$\beta_c/\langle \beta \rangle$ | 0.57 | 0.77 |

Table 5: Best-fit linear regression models for predicting frequency of a cluster in X months time into the future. Terms are listed in the order they were added to the model through forward-selection. The $R^2$Adjusted and Root Mean Squared Error (RMSE) were measured on a testing data set of 5 random time samples over a 25-year period.

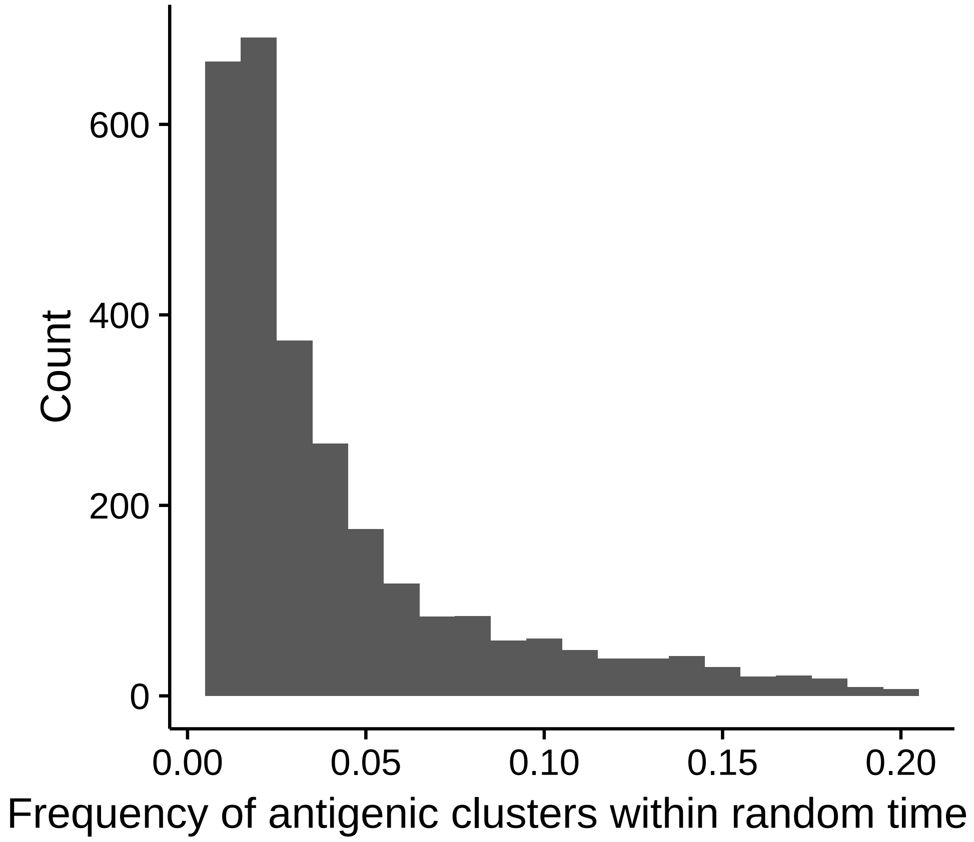


Figure 5: The distribution of the 2846 cluster frequencies from 620 random time samples across the 1500 years of influenza evolution. Clusters that were below 1% relative frequency in the population or those that had already reached our establish criteria were excluded.

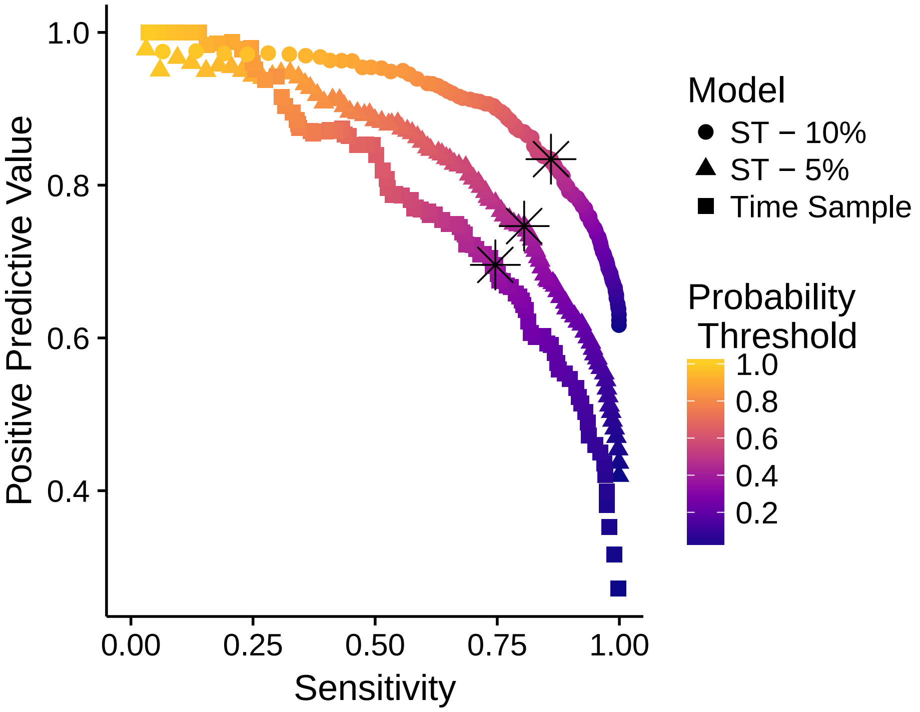


Figure 6: **Comparing the sensitivity and positive predictive value trade-off of two surveillance strategies: 1) surveillance threshold (circles, triangles) and 2) random sampling through time (squares).** Each line highlights the trade off between sensitivity and positive predictive value at different probability thresholds for what constitutes a positive prediction, i.e. a future successful cluster. All models converge in areas with low sensitivity and high positive predictive value, where the model has to predict with probability 0.90 that the cluster will establish in order to classify it as a positive prediction. However, in areas of greater sensitivity, the time sample model consistently under performs surveillance threshold models based on a 5% or higher surveillance threshold. The black stars represent the probability threshold that maximizes the F1 value.

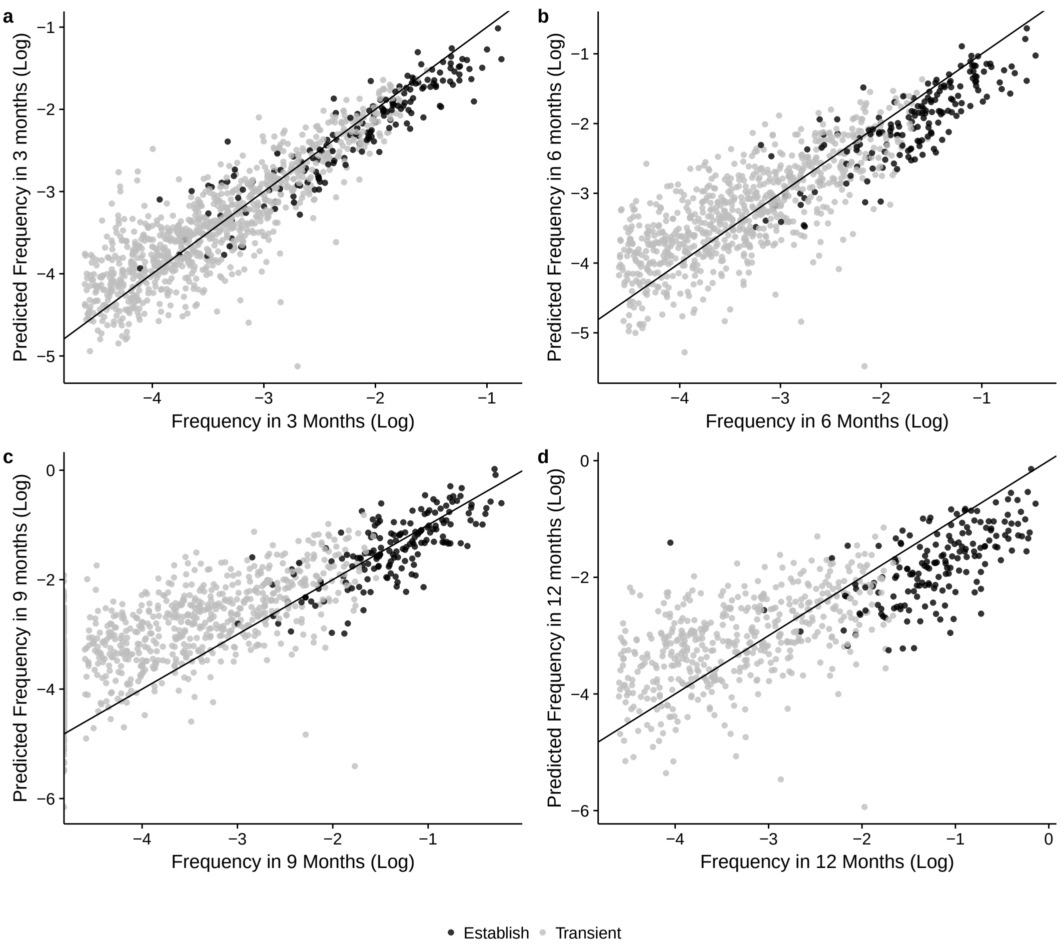


Figure 7: Model predictions of cluster frequency up to a year in advance in three-month increments. Black dots represent clusters that will eventually establish, and grey dots are clusters that will transiently circulate. The black line represents perfect agreement between the actual and predicted log frequencies. The number of clusters present at the time of sampling, but expected to persist in the future, decreases with increasing month-ahead predictions. By 12 months, fewer than half of the starting clusters will still be circulating. As you predict further into the future, the model underestimates future-high frequency circulating clusters, which are usually clusters that will establish.
